# Unleashing the secrets of plant-fungal interactions using a transformation-free confocal staining technique that supports AI-assisted quantitative analysis

**DOI:** 10.1101/2023.10.04.560942

**Authors:** Ashley C. Nelson, Gayan Kariyawasam, Nathan A. Wyatt, Jinling Li, Janine Haueisen, Eva H. Stukenbrock, Pawel Borowicz, Zhaohui Liu, Timothy L. Friesen

## Abstract

Laser scanning confocal microscopy’s ability to generate high-contrast 3D images has become essential to studying plant-fungal interactions. Techniques such as visualization of native fluorescence, fluorescent protein tagging of microbes, GFP/RFP-fusion proteins, and fluorescent labelling of plant and fungal proteins have been widely used to aid in these investigations. Use of fluorescent proteins have several pitfalls including variability of expression *in planta* and the requirement of gene transformation. Here we used the unlabeled pathogens *Parastagonospora nodorum*, *Pyrenophora teres* f. *teres*, and *Cercospora beticola* infecting wheat, barley, and sugar beet respectively, to show the utility of a staining and imaging technique that uses propidium iodide (PI), which stains RNA and DNA, and wheat germ agglutinin labeled with fluorescein isothiocyanate (WGA-FITC), which stains chitin, to visualize fungal colonization of plants. This method relies on the use of KOH to remove the cutin layer of the leaf, increasing its permeability. This permeability allows the staining solution to penetrate and efficiently bind to its targets, resulting in a consistent visualization of cellular structures. We have also used this staining technique in conjunction with machine learning to analyze fungal volume, which indicates the fitness of the pathogen *in planta*, as well as quantifying nuclear breakdown, an early indicator of programmed cell death (PCD). This technique is simple to use, robust, consistent across host species, and can be applied to any plant-fungal interaction. Therefore, this technique can be used to characterize model systems as well as non-model interactions where transformation is not routine.

## Introduction

The progression of understanding plant-fungal interactions has been tightly linked to the development and innovations of microscopy. Light microscopy allows the observation of pathogenic symptom development including the pathogen structures underlying these symptoms (Skipp and Deverall 1972). Both fluorescence and immunofluorescence microscopy are forms of light microscopy that allow visualization of individual fluorophores and specific biomolecules, respectively. After its discovery, green fluorescent proteins (GFP) were expressed in fungal pathogens to track fungal development and used in fusion proteins to allow protein localization in plant cells using fluorescence microscopy (Spellig et al 1996). Immunofluorescence microscopy allowed the use of protein labeling using antibodies to visualize specific molecules in pathogenic fungi (O’Connell et al 2004). These innovations allowed the visualization of specific proteins and locations in both the plant and fungus. However, these tools required fluorescent samples and were subject to photobleaching or signal variability over time and between samples. Electron microscopy (EM) has allowed high resolution imaging of fungal spores, sexual structures, and hyphal plant interactions (Tapio and Photo-Lahdenperä 1991). However, high quality imaging of plant-fungal interactions using EM comes at a high cost with significant training required, and a more laborious process that results in fewer analyzed samples. Confocal microscopy uses multiple lasers, spectral detection, and scanning to separate overlapping emissions from fluorescent proteins to create multispectral images (Bayguinov et al. 2018). These images can be taken through the infected tissue, resulting in improved spatial resolution of the sample. These attributes result in faster imaging of dynamic processes in living cells, detailed morphological analyses of tissues, and protein localization (St. Croix et al. 2005).

Techniques such as visualization of native fluorescence, fluorescent protein tagging of microbes, GFP/RFP-fusion proteins, and fluorescent labelling of plant and fungal proteins have been widely used to study plant-fungal interactions (Todd et al 1999, Jones et al. 2016, Kariyawasam et al. 2022, Dugyala et al. 2015, Haueisen et al. 2019, Redkar et al. 2015, Solanki et al. 2019). These techniques have allowed the detailed visualization of the progression of pathogen colonization within the plant, detailed visualization of virulence factors and the comparison of infection processes when the pathogen is confronted with varying levels of resistance (Jones et al. 2016, Gu and Innes et al. 2012, Todd et al 1999, Jones et al. 2016, Kariyawasam et al. 2022). Using fluorescent proteins has drawbacks including photodegradation (Widengren et al. 1999), variability of expression *in planta (*Lagopodi et al. 2002), difficulty in obtaining high resolution images, and the requirement of gene transformation (Kariyawasam et al. 2022), making these techniques difficult for many agriculturally important non-model interactions. Additionally, doing large scale comparisons of multiple interactions is not feasible due to transformation being labor intensive, even in model systems.

In this study, we set out to develop a straightforward confocal staining and imaging technique that yields consistent results between samples, with improved visualization for both the plant and fungal pathogen using contrasting fluorescent stains that eliminate the need for gene transformation. As proof of concept, we used the non-fluorescently transfected foliar pathogens *Parastagonospora nodorum* (septoria nodorum blotch), *Pyrenophora teres* f. *teres* (net form net blotch), and *Cercospora beticola* (Cercospora leaf spot) of wheat, barley, and sugar beet respectively, to show the efficacy of this technique. The staining technique was subsequently coupled with machine learning to analyze fungal volume, an indicator of pathogen fitness *in planta*, and nuclear breakdown, a classical early indicator of programmed cell death (PCD). Machine learning volume analysis propels fungal volume detection methods forward, as other methods, like qPCR, are laborious and only serve as a proxy to estimate the fungal biomass, while this method shows the actual estimate of fungal volume (Feckler et al. 2017, Kumar et al. 2015). This technique is simple, robust, durable, consistent, and can be used to characterize model and non-model plant-fungal interactions without transformation, making it useful to anyone using confocal microscopy to characterize plant-fungal interactions.

## Methodology

### Sample Collection

#### Parastagonospora nodorum - wheat interaction

The wheat line Grandin was grown for 14 days, or until the secondary leaf was fully expanded (Figure 1A). Attached secondary leaves were taped down onto a high-density polyethylene (HDPE) board and inoculated with fresh spores of the *P. nodorum* isolate, LDN03 Sn-4 (hereafter referred to as Sn4), that is virulent on Grandin (Figure 1B). Inoculation and spore preparation was done as described in Liu et al. (2004). Briefly, *P. nodorum* cultures were grown by placing a dry agar plug containing Sn4 onto V8-PDA (150mL of V8 juice, 10g of Difco potato dextrose agar, 3g of CaCO_3_, and 10g of agar in 1,000mL of water) and allowed to rehydrate for one hour, then streaked across the plate to distribute the fungus. The plate was incubated at room temperature under continuous light for six days, or until pycnidia emerged. Each plate was flooded with sterile distilled water and pycnidia were agitated with a sterile pipette tip to stimulate the release of pycnidiospores. The spores were harvested, and the spore solution was adjusted to a concentration of 1 x 10^6^ spores/mL. Two drops of the surfactant Tween20 (J.T. Baker Chemical Co.) were added per 100mL of spore suspension. Inoculation of attached leaves was done using a pneumatic paint sprayer (Husky) and inoculated until run-off of the spore solution was observed (Liu et al. 2004). After inoculation, the plants were kept in a mist chamber for 24 hours under continuous light and 100% relative humidity at ∼ 21 °C to facilitate spore germination and fungal penetration (Figure 1C). After 24 hours, the plants were moved to a growth chamber under a 12-hour photoperiod at 21 °C for the remainder of the experiment (Figure 1D). Three leaf tissue samples were collected at 12, 24, 48, 72, and 96 hours post inoculation (hpi) (Figure 1E). After collection, each set of leaves at each timepoint was photographed using a Canon EOS 70D attached to a copy stand and then prepared for staining. This was repeated twice more, resulting in three biological replicates.

**Fig. 1.**
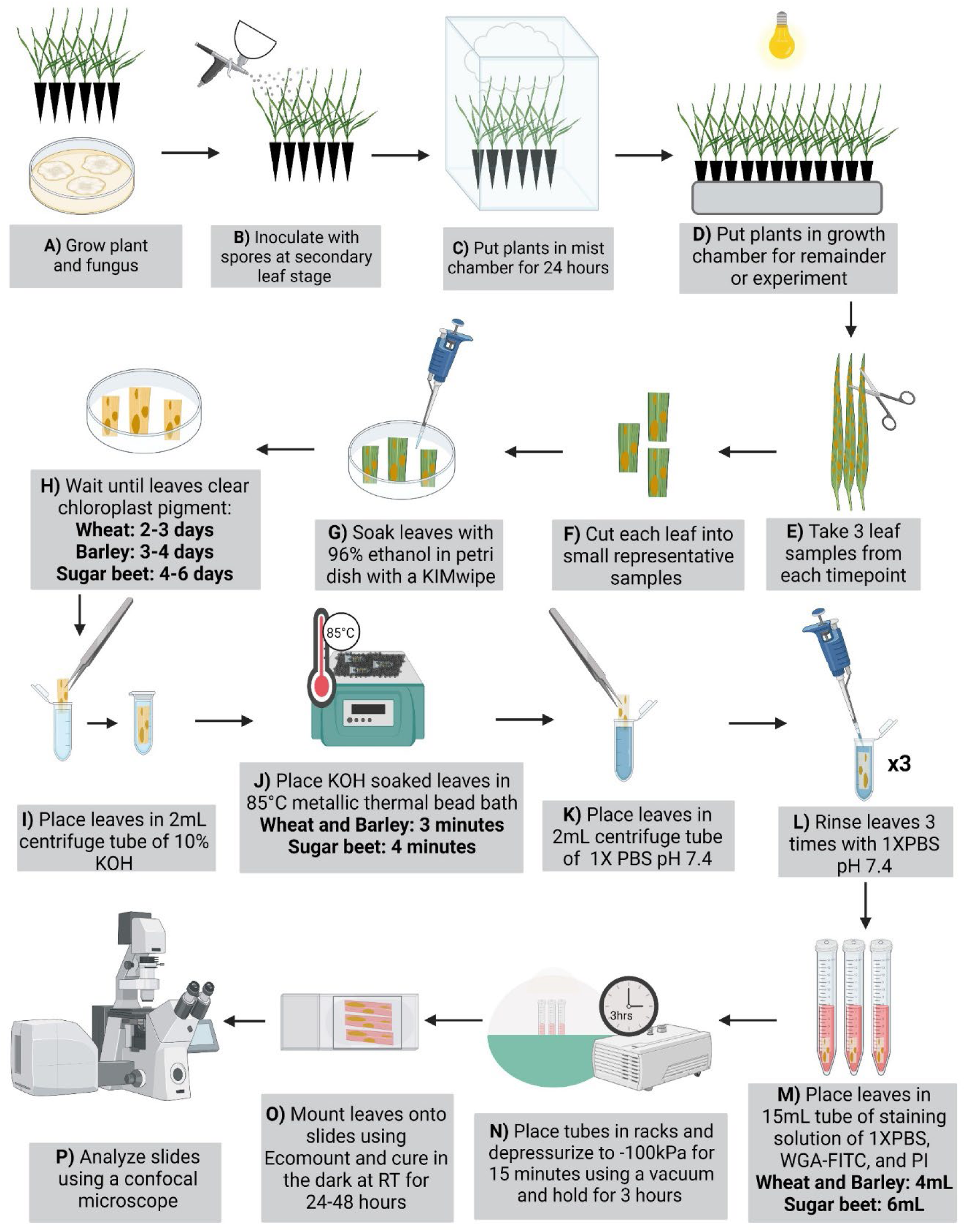
Process of foliar sample collection and staining using WGA-FITC and PI.

#### Pyrenophora teres f. teres – barley interaction

The barley line Rika was grown as stated above for wheat but inoculated with the *P. teres* f*. teres* isolate #5*VR1*/*VR2* that is highly virulent on Rika (Shjerve et al. 2014) (Figure 1A). Spore growth and inoculation were performed as described in Koladia et al. (2017). Briefly, *P. teres* f. *teres* isolate #5*VR1*/*VR2* was grown on plates containing V8-PDA media in the dark at room temperature for five to seven days. Subsequently, plates were placed under continuous light at room temperature for 24 hours and moved back into the dark for 24 hours at 15°C. The plates were flooded with 100mL of sterilized distilled water and scraped using an inoculation loop, resulting in conidia release. The spore solution was then diluted to 2,000 spores/mL with 1 drop of Tween20 added for every 50mL of spore solution. Inoculation of the spore suspension on barley was done as described for wheat (Figure 1B). Plants were kept in the growth chamber and followed the conditions mentioned above in wheat and samples were collected at the same timepoints (Figure 1A-G). One replication was completed.

#### *Cercospora beticola* - sugar beet interaction

The susceptible sugar beet cultivar Crystal 093 (Beta seed, Frankfurt am Main, Germany) was inoculated with the virulent *C*. *beticola* isolate 309-10 collected from southern Minnesota, USA. Isolate 309-10 was cultured on CV8 agar (177mL V8 juice, 3 grams CaCO_3_, 16 grams agar and 823mL H_2_O per 1 liter of media) plates for 14 days at 30 °C with a 16-hour photoperiod. After 14 days, *C. beticola* cultures were scraped with a sterile glass slide to remove mycelial tissue from the plate. Cultures were allowed to air dry before being incubated at room temperature in 24 hour light conditions. After 48 hours, conidia were harvested using 10mL of sterile water. The spore suspension was adjusted to a concentration of 2,000 spores/ml and evenly sprayed on leaves of 5-week-old Crystal 093 sugar beet plants. Inoculated plants were kept in a humidity chamber at 27 °C and 90% humidity for five days after which plants were kept at 22 °C with a 16hr photoperiod. Leaves were sampled for staining at 5, 7, 10, and 14 days post inoculation (dpi). One replication was completed.

### Staining

The staining process was validated on the *P. nodorum* -wheat, *P. teres* f. *teres*-barley, and *C. beticola-*sugar beet interactions. Leaf tissue samples from each inoculated plant from each timepoint were harvested and cut into 3cm × 0.5cm sections (Figure 1F) and cleared in a parafilm sealed petri dish using a KIMwipe (Kimberly Clark, TX, USA) soaked in 96% ethanol to remove chlorophyll pigments (Figure 1G). The KIMwipe was continually soaked with 96% ethanol until leaves were cleared of all chlorophyll pigment. Leaves were considered cleared when no green was visible resulting in a pale yellow/tan color (Figure 1H). Once leaves were cleared (wheat 2-3 days, barley 3-4 days, and sugar beet 4-6 days), each individual leaf was immersed in 10% KOH (w/v) in a 2mL centrifuge tube and placed in an 85°C metallic thermal bead bath (Thermolyne dri-bath with Lab Armor beads) for three minutes (four minutes for sugar beet) (Figure 1I and 1J). KOH is effective at removing the cutin layer of the leaf, making the leaf more permeable and allowing penetration of the stain into the leaf. Immediately after KOH submersion, each leaf was placed in an individual 2mL centrifuge tube and rinsed three times with a 1× phosphate buffer solution (PBS) pH 7.4. This was diluted from 10× PBS pH 7.4 that was made according to the Cold Spring Harbor protocol (25.6g Na_2_HPO_4_, 80g NaCl, 2g KCl, and 2g KH_2_PO_4_ per 1 liter of H_2_O, Cold Spring Harb. Protoc. 2007) (Figure 1K and 1L). Each leaf was submerged into an individual 15mL conical centrifuge tube containing 4mL of staining solution for wheat and barley and 6mL of staining solution for sugar beet (Figure 1M). One milliliter of staining solution consisted of 1mL 1×PBS, 10µg/mL of wheat germ agglutinin labeled with fluorescein isothiocyanate (WGA-FITC, Sigma Aldrich L4895) and 20µg/mL propidium iodide (PI, Sigma Aldrich P4170). WGA-FITC binds chitin in the fungal cell walls, while PI binds DNA and RNA. The staining solution was kept away from the light by wrapping each tube with aluminum foil and the top was covered with parafilm. A hole was placed in the parafilm at the top of each tube to prevent the solution from bubbling. The 15mL tubes with the staining solution and leaves were placed in racks inside a vacuum desiccator and depressurized to −100kPa for 15 minutes using a two-stage rotary vane vacuum pump (Edwards 8.4 CDM RV12) and held at constant pressure for three hours (Figure 1N). After depressurization, the three leaf tissue samples from each treatment at each timepoint were mounted on a microscope slide using EcoMount (BioCare Medical, USA) (Figure 1O). Leaves were oriented with the adaxial surface facing up and a 22×40mm #1.5 coverslip (VWR Microscope Cover Glass #1,5) was placed on top, avoiding air bubbles. The slides were left to cure in the dark for 24-48 hours at room temperature then placed into a slide box and stored at 4°C until they were ready for confocal analysis (Figure 1P).

### Confocal Laser Scanning Microscopy analysis

All slides for all three interactions were observed under a Zeiss LSM700 laser scanning confocal microscope (Zeiss, Germany) using both 20× 0.8 NA dry, and 40× 1.45 NA oil immersion objectives. Three different channels were used, a red channel (Ex555/Em 630nm) captured the fluorescence emitted by propidium iodide (PI), which binds nucleic acid (DNA and RNA) in the nucleus and cytoplasm (Suzuki et al. 1997). A green channel (Ex488nm/Em520nm) captured fluorescence emitted by wheat germ agglutinin labeled with fluorescein isothiocyanate (WGA-FITC), which binds chitin in the fungal cell wall, and a blue calcofluor white (CaFW) channel (Ex366nm/Em420nm) captured autofluorescence of plant leaf/cellular structures. The infection process was observed in different tissue layers within the leaf using 3D Z-stack images. Two technical replicates of each biological replicate were completed, noting observations, and capturing representative images. At least six representative images were taken at each timepoint for each treatment at both 20× and 40× magnification.

### Image analysis

Images for each of the three interactions were analyzed using the Imaris 10.0.0 software (Oxford Instruments). Confocal microscopy images (.lsm files) from each timepoint and treatment were uploaded to Imaris (Oxford Instruments), channel emission was equalized for blue, red and green colors. Images were captured from the 3D view, then cropped to 102µm × 102µm × 46µm (X × Y × Z) for final use.

### Volume analysis

Volume analysis was performed for each of the three interactions by creating a surface in Imaris v.10.0.0 of the fungal growth. The “segment only a region of interest”, “classify surfaces”, and “object-object statistics” options were unchecked in the primary window. After selecting the next step, the “smooth option” was unchecked and selection for “labkit for pixel classification” and “all channels’” options were selected. Selecting “next” brings you to the LABKIT Fiji plugin. The LABKIT Fiji plugin uses pixel machine learning that is trained by identifying foreground (fungus) and background (plant cells) (Arzt et al. 2022). The LABKIT plugin is compatible with both the Imaris and ImageJ software and can be installed using the GitHub page: https://github.com/juglab/labkit-ui. Three to five rounds of training were done to create individual algorithms for *P. nodorum*, *P. teres* f. *teres*, and *C. beticola* and saved as a classifier file in Fiji. These algorithms were run on each host-pathogen combination at each timepoint, then exported back to Imaris as surface data. After the surface was created, and voxel noise was minimized, statistics were selected via the statistics tab, and exported to an excel spreadsheet containing a sum of fungal volume (μm^3^) column (Figure 2). The summed volume statistic was taken from each image, analyzed per timepoint, averaged among the other volumes recorded and then standard error of means were calculated. The average volume and standard deviation for each timepoint was then used to create a graph (Figure 4).

**Fig. 2.**
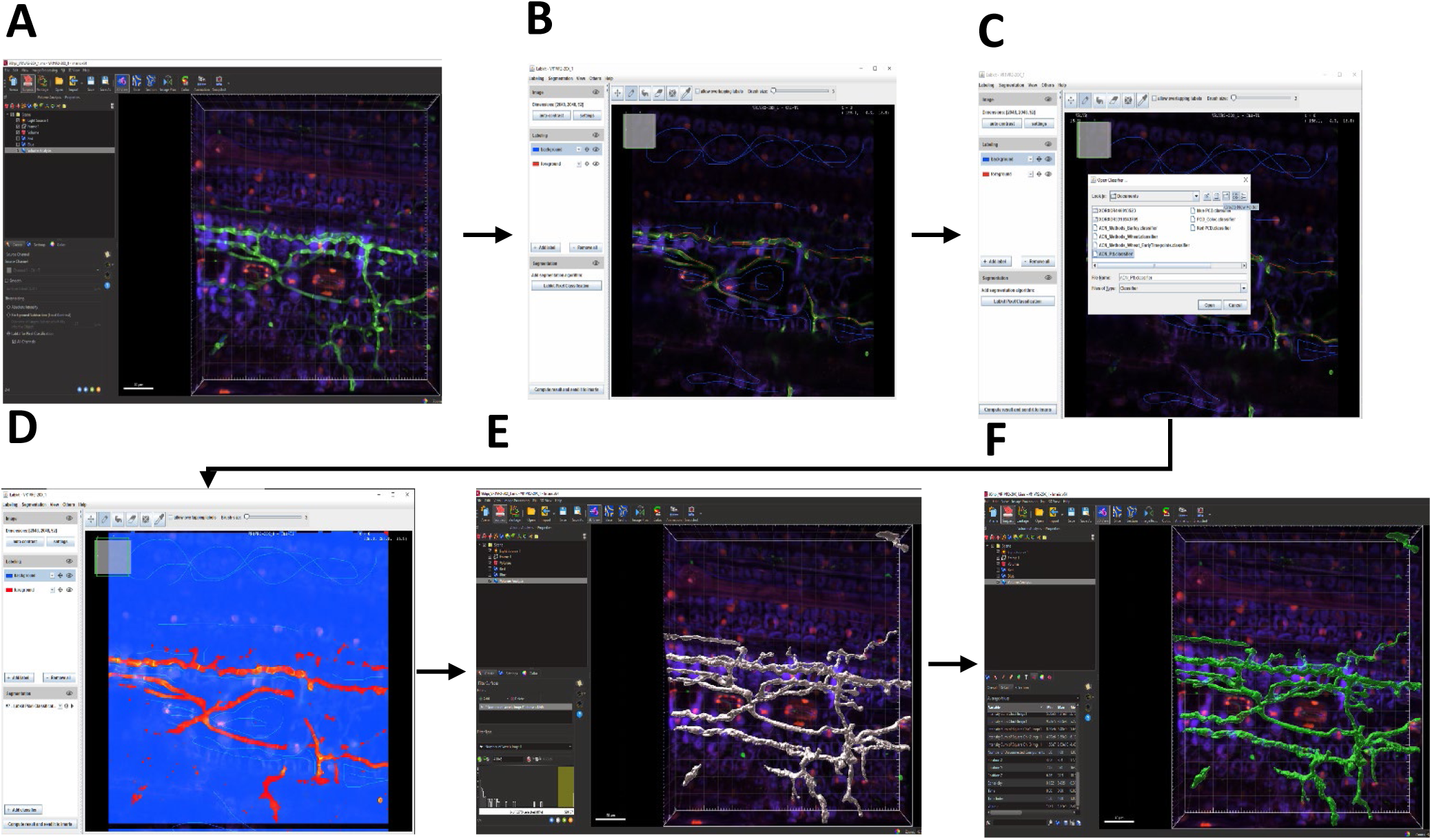
Volume Analysis Using Imaris 10.0.0. **A**, Starting on Imaris 10.0.0 with a confocal image, a surface is created and the option for using LABKIT with pixel classification is selected. **B**, This brings you to a Fiji plug in where the drawing mode is selected. The volume of interest, in this case the fungus (green) is drawn in with red which represents the foreground and the plant autofluorescence and DNA (blue and red respectively) as well as stomata are drawn in with blue which represents the background. **C**, From this the pixel classification machine learning is trained and an algorithm can be saved. In this case we are using the Ptt.classifier algorithm to characterize *P. teres* f. *teres* on barley. **D**, Once this algorithm is formulated and ran it results in the blue and red pixelated image that is observed above outlining the fungus. **E** & **F**, This is then computed, sent to Imaris where the voxel number is specified and the surface representing fungal volume is calculated and placed into the 3D image.

### Loss of nuclear volume as an indicator of programmed cell death

#### *P. nodorum*-wheat interaction

Programmed cell death (PCD) analysis was performed by visualizing the nuclear degradation across all timepoints in the *P. nodorum*-wheat system. Nuclear degradation was assessed by creating a surface in Imaris v.10.0.0 using the LABKIT Fiji plugin as described in the volume analysis above. The algorithm was made using eight rounds of training with nuclei as the foreground and other plant and fungal components as the background. Once the surface was created it was named and volume statistics of the nuclei were extracted as described above. The summed volume statistic from each image at each timepoint was averaged and standard error of mean was calculated, and used to make a graph (Figure 4).

## Results

### Staining method using three different plant-fungal interactions

To confirm that this method could be used for visualizing pathogens associated with different host types, we used the *Pyrenophora teres* f. *teres* – barley, *Parastagonospora nodorum* – wheat, and *Cercospora beticola* – sugar beet interactions. Host species were inoculated with virulent isolates of their respective pathogens, stained, and visualized using the Zeiss LSM700 laser scanning confocal microscope. In these images, the fungal cell wall component (chitin) was stained using WGA-FITC and were visualized in green, the nuclei were stained using PI and were visualized in red, and the plant cell autofluorescence was visualized in blue.

Micrographs of the *P. teres* f. *teres*-barley interaction and *P. nodorum* – wheat interaction were taken at 12, 24, 48, 72 and 96 hpi (Figure 3A) and the *C. beticola*-sugar beet interaction was analyzed at 5, 7, 10 and 14 days post-inoculation (dpi) (Figure 3B). In those images, contrast color differentiation was observed between fungal hyphae (green), plant nuclei (red), and other plant structures (blue) (Figure 3). The high resolution and detailed images of the staining allowed us to follow the morphology of the fungal growth patterns from spore germination on the surface of the leaf through extensive mesophyll colonization. We can also observe small details such as putative feeding structures, septa, and hyphal growth orientation (Figure 3). By staining both the fungus and plant in one protocol and accounting for autofluorescence, we are able to clearly differentiate the fungal activity in the plant. Given the differentiation and detail in each image, a volume analysis can be conducted to accurately estimate fungal volume and nuclear degradation, an early hallmark of PCD. The use of wheat, barley, and sugar beet demonstrate that the staining method is effective for both monocot and dicot plant-fungal interactions, indicating that this staining protocol will be useful for a broad range of foliar fungal interactions and may also be extended to other plant tissues with small protocol modifications.

**Fig. 3.**
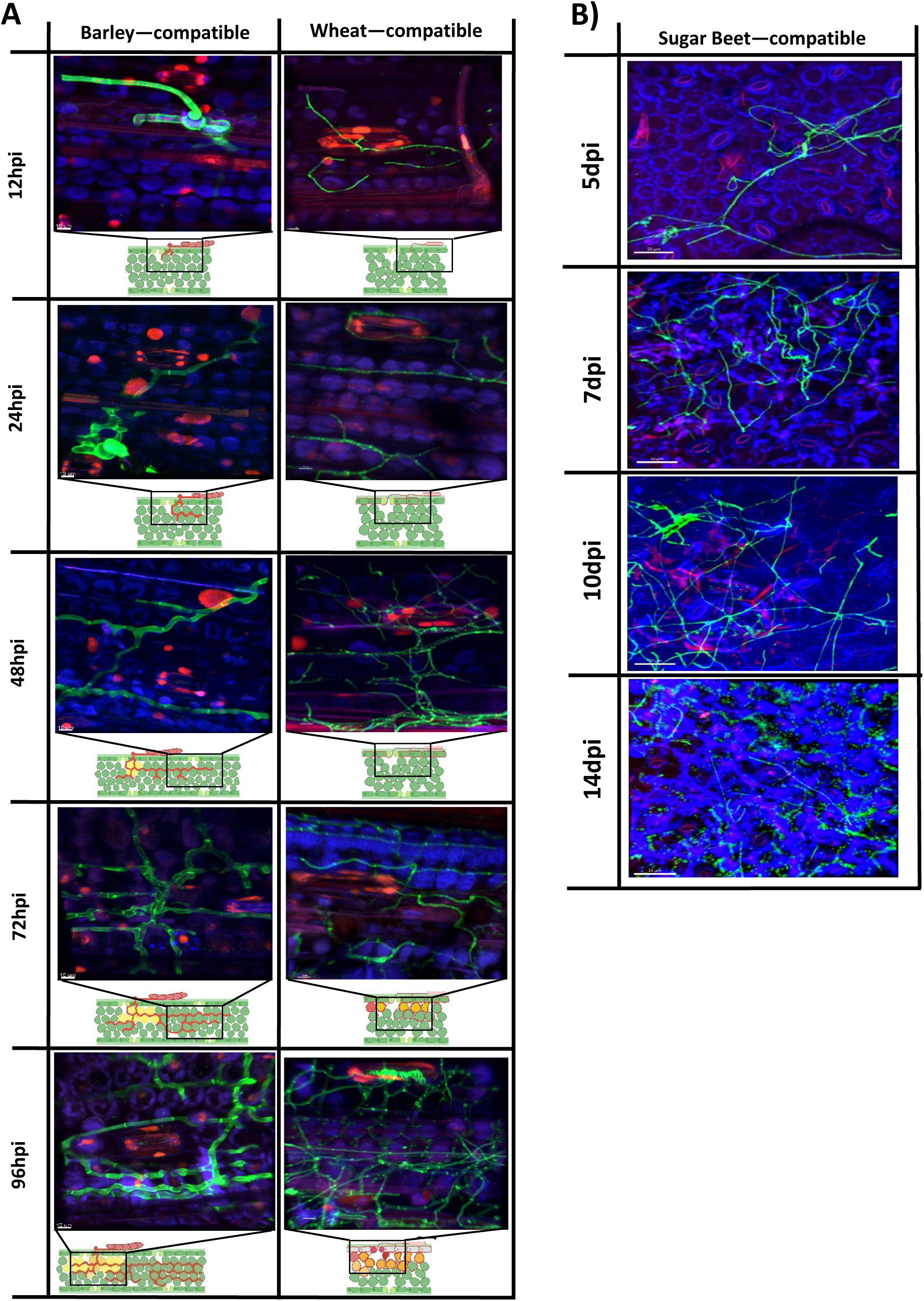
Disease progression of compatible interaction. **A**, Micrographs of disease progression for the compatible barley *– Pyrenophora teres* f. *teres* interaction and the wheat – *Parastagonospora nodorum* interaction at 4, 12, 24, 48, 72 and 96 hours post inoculations (hpi). In the barley-*P. teres* f. *teres* interaction an intracellular vesicle is established in the epidermal cell and the start of mesophyll colonization has begun. By 24hpi mesophyll colonization has started to reach further into the leaf. At 48hpi the fungal hyphae have established themselves in the mesophyll layer focusing on growing parallel to the epidermal layer. By 72hpi the pathogen had established multiple layers of parallel growth in the leaf mesophyll layer and perpendicular branching is taking place. At 96hpi the pathogen has continued to colonize any free apoplastic space and encircles mesophyll cells. In the wheat*-P. nodorum* reaction spore germination and pathogen penetration into the leaf takes place at 12hpi. At 24hpi intercellular colonization of the epidermal cell takes place. From the epidermal colonization, intercellular colonization begins in the top mesophyll layer at 48hpi. At 72hpi the mesophyll colonization goes deeper into the leaf eventually progressing deep through the leaf by 96hpi. **B**, Micrographs of disease progression for the compatible reaction sugar beet – *Cercospora beticola* at 5, 7, 10 and 14 days post inoculation (dpi). Fungal hyphae are displayed in green while blue shows plant autofluorescence and red corresponds to DNA. At 5dpi spore germination takes place on the surface of the leaf with penetration taking pace by 7dpi. At 10dpi colonization of the epidermal takes place and by 14dpi extensive mesophyll colonization can be observed. Cartoon figures were created using the web-based application BioRender (BioRender.com). Yellow, orange and red mesophyll cells in the cartoons represent different stages of programmed cell death (PCD) inside the leaf. Cells in early stages of PCD are yellow, red cells are collapsed cells and orange cells are in the middle of the PCD process

### Volume Analysis

#### P. nodorum-wheat

In the compatible reaction, the fungal volume at 12hpi, 6×10^4^ μm^3^, was followed by a slight increase at 24hpi to an average of 7×10^4^ μm^3^ (Figure 4). This increase represents spore germination and the initiation of intercellular epidermal colonization observed at these timepoints (Figure 3A). At 48hpi, the fungal volume significantly increased, using standard error of mean at a *p* value of 0.05, to 2.4×10^5^ μm^3^ (Figure 4). In Figure 3A, extensive intercellular epidermal colonization seen at 48hpi explains the increased fungal volume. At 72hpi, there was a relatively small increase in total fungal volume (averaging 2.7×10^5^ μm^3^) (Figure 4) marked by the initiation of mesophyll colonization in addition to the continued progression of extensive intercellular colonization of the epidermal layer (Figure 3A). At 96hpi a fungal volume of 4.3×10^5^ μm^3^ (Figure 4) was observed and is reflected by extensive colonization of both the epidermal and mesophyll cells showing near complete colonization (Figure 3A). These results indicate that this staining technique produces consistent images that can be used for accurate fungal volume analysis.

**Fig. 4.**
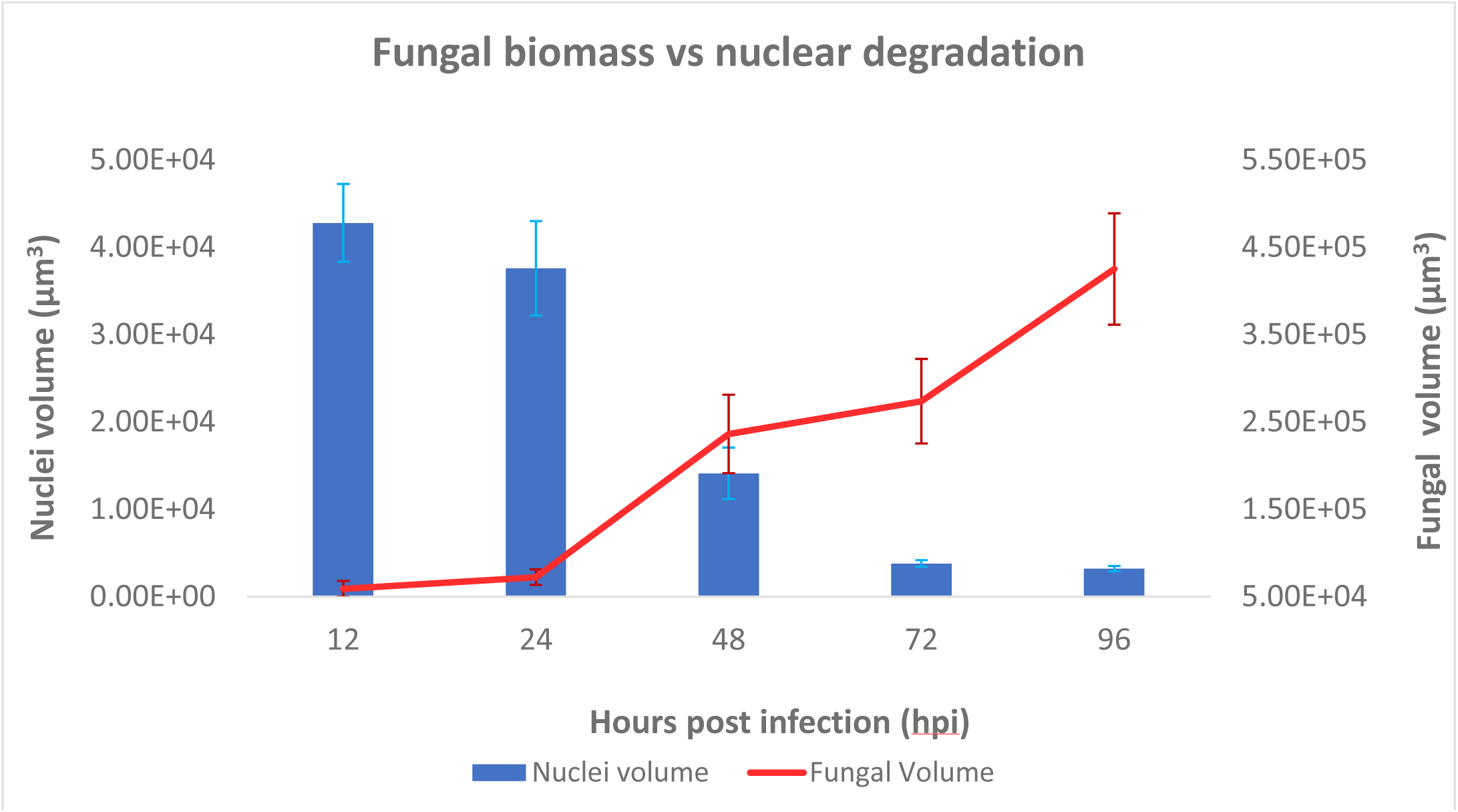
Fungal volume vs nuclear degradation (PCD) Fungal volume of *Parastagonospora nodorum* on wheat and plant nuclear degradation (used to assess PCD) at the timepoints 12, 24, 48, 72, and 96hpi. Fungal volume is in red and begins at 5.00E+04 and has a range up to 5.50E+05 while nuclei volume is in blue and has a rang up to 5.00E+04. A decrease in nuclei volume is representative of programmed cell death (PCD). Error bars represent the standard error of mean (SEM) from three replications for each timepoint. The fungal volume of *P. nodorum* increased after 24 hpi while the nuclear volume (PCD indicator) decreased after 24 hpi.

#### *P. teres* f. *teres*-barley and *C. beticola*-sugar beet

Volume analysis for both the *P. teres* f. *teres*-barley and the *C. beticola*-sugar beet interactions was done at 12, 24, 48, 72 and 96 hours post inoculation (hpi) and 5, 7, 10 and 14 days post inoculation (dpi), respectively to validate that the process used for the *P. nodorum*-wheat volume analysis could be applied to these two systems. It was found that algorithms developed for each host pathogen complex (described previously) worked effectively and volume analysis reflected the visual observations represented in Figure 3 and therefore can be used in both monocot and dicot plant interactions (Supplementary Figure 1).

### Nuclear Degradation

#### P. nodorum-wheat

Volume analysis on plant nuclei was done using the same 20x magnification images analyzed for fungal volume, resulting in six representative images at each timepoint. In the compatible reaction, we observed a progressive decrease of plant nuclei volume across timepoints. Nuclear degradation is known to be an early hallmark of PCD and therefore can be used as a proxy for quantifying PCD (Figure 4). There was no significant difference of nuclear volume between 12 and 24hpi, which were around 4×10^4^ μm^3^ (Figure 4). A significant decrease in nuclear volume(about a 60% reduction in volume) was observed at 48hpi (1.4×10^4^ μm^3^). This is followed by another significant decrease at 72hpi with around an 80% reduction to 3×10^4^ μm^3^. This low nuclear volume stays stable through the 96hpi timepoint. These results indicate that using volume analysis to evaluate nuclear volume can be used to track PCD.

## Discussion

We have developed a staining and imaging technique that allows for the visualization and analysis of multiple plant-fungal interactions, while eliminating the need for transformation of fungal strains expressing fluorescent proteins. The staining and imaging technique presented here uses commonly available reagents that permit high resolution images of fungal growth, resulting in accurate observations and quantification through volume analysis. By adapting this technique from previously established staining protocols (Haueisen et al. 2019) that used wheat germ agglutinin labeled with fluorescein isothiocyanate (WGA-FITC) and propidium iodide (PI) we were able to make changes to improve its resolution, replicability, and adaptability across numerous pathosystems. Crucial aspects such as the reagents (WGA-FITC, PI and 1XPBS pH 7.4), clearing leaves using 96% ethanol and 10% KOH heat treatment were retained while depressurization duration, pressure and equipment was adapted alongside the use of three lasers in confocal analysis, and the establishment of AI assisted quantitative analysis sets this technique apart. This highly accurate yet relatively simple staining technique is effective in both monocot and dicot plant-fungal systems (necrotrophs and hemi-biotrophs), showing its broad utility in plant-fungal interactions.

The success of this staining process hinges on unpigmented leaves, increased leaf permeability, both plant and fungal staining reagents, and vacuum depressurization using standard laboratory equipment. Removing the chlorophyll pigments from the leaves using 96% ethanol, allows more effective uptake of the staining reagents and prevents interference from chlorophyll auto-fluorescence. Clearing time will vary depending on the leaf type, as larger and thicker leaves will take more time to clear. When using thicker leaves such as sugar beet, the original ethanol soaked KIMwipe may need to be replaced, as it can soak up the chlorophyll pigment relatively quickly. Leaves can also be stored in ethanol for longer periods of time before microscopy, we found that storing the cleared leaves in ethanol for a week or less gave the most consistent results. Using leaves stored in ethanol for two weeks resulted in less consistent visualization of the fungus.

Increasing the permeability of the leaves using KOH allows the staining solution to penetrate deep into the plant and fungal cells. KOH is an alkali which digests components surrounding the fungal cell wall allowing clear visualization of the hyphae (Cheesbrough 2005). Obtaining the correct balance of permeability is a delicate process, and we found that heat and a 10% KOH treatment was highly effective. The amount of time the leaves were exposed to this treatment was critical, with under treatment (shorter amount of time), resulting in a stiff leaf with an intact or partially intact cutin layer, prohibiting the staining solution from entering the plant cells. KOH undertreatment also resulted in lower quality and blurred staining of fungal hyphae by WGA-FITC (Hood and Shew 1996). However, excessive exposure to this treatment can result in the deterioration of the leaf, leading to total leaf collapse and the inability to treat the leaf in subsequent steps of this protocol. KOH treatment timing is dependent on the characteristics of the plant leaf; Smaller and thinner leaves, such as wheat, require less treatment time compared to larger, thicker leaves such as sugar beet.

The staining reagents are important in highlighting the fungal growth through the visualization of the mycelium (chitin) stained with WGA-FITC and plant nuclei (RNA/DNA) stained with PI. The concentrations of 10 µg/mL of WGA-FITC and 20 µg/mL of PI were optimal across all three host-pathogen systems. We tested increased concentrations of WGA-FITC (20 µg/mL) and PI (40 µg/mL) which resulted in oversaturated images. Optimal depressurization was consistent across all three host-pathogen systems, with 15 minutes of active depressurization of −100kPa and three hours of holding depressurization being the most effective. We found that leaves with 5 and 10 minutes of active depressurization didn’t contain enough stain in the mesophyll layer compared to leaves with 15 minutes of active depressurization. Active depressurization longer than 15 minutes had the same amount of mesophyll layer staining as the 15-minute treatment, therefore, we concluded that 15 minutes was optimal. We also attempted active depressurization rotated with holding depressurization every five minutes for one hour. This resulted in inconsistent staining of the cellular levels of the leaf and therefore we focused on a constant pressure. We tested one hour, three hour, and overnight holding of depressurization. Depressurization for one hour did not provide full and consistent staining of all leaves while overnight and three hours of depressurization had similar staining, therefore we used the three hours of constant pressure for best results.

If adapting this protocol for another system, the most critical elements are the optimization of the clearing time and the 10% KOH treatment time. Optimizing these areas results in the correct amount of staining uptake, resulting in the best differentiation of plant and fungal tissue. For clearing and KOH treatment time, the thickness and surface area of the leaves collected need to be considered. When testing KOH treatment times for wheat, we used two-minute, three-minute and four-minute treatments since wheat secondary leaves are thin and narrow. We found that the two-minute treatment left the leaf cutin layer partially intact while the four-minute treatment resulted in leaf collapse that couldn’t be transferred to other steps in the protocol. The three-minute interval was optimal because it provided a permeable but structurally stable leaf that could consistently take up the staining solution. For testing the KOH treatment on barley, we considered that it is thicker and wider than the wheat leaves and therefore used three minutes as the first timepoint, followed by four minutes and five minutes. We found that at five minutes the barley leaf had completely collapsed, while the three minute and four-minute barley leaves were similar in permeability and leaf flexibility. After laser scanning confocal microscope evaluation of the stained leaves, both the three- and four-minute time points had similar staining depth and coverage. Given this, we moved forward with the three-minute treatment time for barley.

Moving to sugar beet, the dicot has much broader and thicker leaves than barley or wheat. Three-, four-, five- and six-minute treatment times were compared. The three-minute timepoint left a partially intact cutin layer on the leaf that would not be effective for stain uptake while the six-minute treatment resulted in leaf collapse. Both the four- and five-minute timepoint fully removed the leaf cutin layer, with the five minute treatment leaf being slightly more flexible than the four minute treatment. When compared at the microscope after staining, we found similar staining throughout the leaves for both the four- and five-minute treatments. Given this, we moved forward with the four-minute treatment time for sugar beet. The staining technique presented here, complemented by mounting the leaves in EcoMount, contributed to the durability of this procedure. EcoMount dries relatively quickly and is an optically clear mounting medium that preserves the samples for at least one year when stored properly.

The technique presented here has several advantages over other visualization techniques. Using fluorescent protein tagging, the fluorescent protein intensity can decrease over time and become unusable less than a year after transformation. Our technique was developed to be simple, using only readily available reagents, and requiring common lab equipment. The staining protocol is consistent and repeatable with minor adjustments using three different host-pathogen systems with three different leaf morphologies. This was also serendipitously tested by multiple lab members, resulting in similar success even with some variability in technique.

Confocal microscopy is an essential part of 3D images where multiple lasers are required to excite the WGA-FITC and PI resulting in the emitting of contrasting colors. Using a combination of staining reagents that have different wavelengths is critical in differentiating plant and fungal structures including plant nuclei and fungal mycelia. However, some fungal spores (e.g., *P. teres* f. *teres)* have a gelatinous layer and WGA-FITC staining will not penetrate enough to stain the chitin in the cell wall, and therefore are not easily visualized. Accounting for plant autofluorescence is equally important when analyzing and distinguishing fungal growth, which was done by using a blue, CaFW, channel and clearing leaves to eliminate chlorophyll autofluorescence.

Images taken with a 20x objective provided a broader view while 40x objective (oil immersion) provided more detail, and when used together, provide a comprehensive view of the interaction. An example of this can be seen when comparing the *P. nodorum*-wheat and *P. teres* f. *teres*-barley pathosystems. *P. nodorum* is a spotting pathogen and therefore colonization takes place in focused oblong areas in the leaf. This interaction is best visualized using a higher magnification objective that can better visualize highly localized colonization that goes deep into the leaf. As for *P. teres* f*. teres*, its colonization is net-like and typically colonizes lengthwise toward the tip and base of the leaf without going as deep into the leaf relative to *P. nodorum*. This parallel expansion is best captured by an objective with a wider field of view, typically a lower magnification. The 20x objective is preferred for volume analysis, as it captures a greater area of the leaf to give a broader look into fungal colonization based on a volume analysis. By analyzing volume over a larger area, more accurate data can be collected. However, it is important for each researcher to understand their fungus (pathogenic, endophytic, symbiotic, etc.) before imaging to decide which objective should be used to provide images specific to their research objectives. In many pathosystems there are situations where both lower and higher magnifications are useful and we recommend taking micrographs at multiple magnification. For example, *P. teres* f. *teres* focuses on growing lengthwise through the leaf mesophyll layer and this behavior is best captured by a lower magnification. However, it also forms intricate structures on the surface and the epidermal layers of the leaf where higher magnifications are useful to capture more detail.

Image analysis can be done on any software that can analyze and edit .lsm files obtained by a confocal microscope, but in this study Imaris v.10.0.0 was used since it has all the modules for analyses, including machine learning. Once captured, images can be cleaned, edited, and sized to preference for analysis and publication. The Imaris software (versions 9.0.0 and newer) plugin tool LABKIT was used to analyze fungal cell and plant cell nuclear volume. This machine learning tool proved powerful for accurate and efficient volume analysis but may not be available if using other software packages. It has been shown to be compatible with both Imaris and ImageJ, and more information can be found in Arzt et al. (2022). We are not endorsing or advocating for the purchase of Imaris but rather just describing useful and innovative tools to assess and quantify pathogen colonization patterns in plants.

This technology was used to analyze plant nuclei volume over time to assess programmed cell death (PCD) in plant leaves. Nuclear dismantling and fragmentation are an early hallmark of plant PCD, and a decrease in nuclear volume overtime is a good proxy to monitor the PCD process (Danon et al. 2000, Dominguez and Cejudo 2015). We created a machine learning algorithm using the LABKIT Fiji interface, that went through multiple rounds of training and redevelopment. A challenge with assessing nuclei in an image is identifying faint shades of the nuclei (red) in an image while also differentiating it from stomata. Precise training and identifying of the foreground are essential for obtaining accurate results.

This staining and imaging technique uses common staining reagents but combines them in an innovative way to ensure detailed and high-quality images while being cost effective, and time saving. By pairing both the staining and imaging techniques, we have provided a protocol that improves colonization assessment, and statistical analysis. Using this protocol allows quick sample and imaging processing when using a larger sample set. We have shown this method to be effective using three different foliar pathogens on three different host plants, including one dicot and two monocots. However, this method is potentially useful for any plant-fungal interaction. Each step in this process is robust, replicable and can withstand small variation in technique making it easily adaptable.

## Acknowledgments

This work was supported [in part] by the U.S. Department of Agriculture, Agricultural Research Service through project 3060-22000-051-00D.

Mention of trade names or commercial products in this publication is solely for the purpose of providing specific information and does not imply recommendation or endorsement by the U.S. Department of Agriculture. USDA is an equal opportunity provider and employer.

The authors would like to thank Danielle Holmes for technical assistance, Dr. Amber English at Imaris for technical software support and Beta seed for providing Crystal 093 seed

## Author contributions

TLF, ACN, and GKK designed the research; ACN and GKK developed the modified protocol on wheat; ACN developed the protocol on barley and sugar beet; ACN performed the confocal imaging and volume analysis for all interactions; NAW performed sugar beet inoculation; JL performed barley inoculation; PB provided expertise in microscopy; ZL provided oversight; TLF and ACN wrote and organized the manuscript. All authors edited and approved the manuscript.

## Notes

Funding: National Institute of Food and Agriculture Grant Number: 2016-67013-24813 and 2018-67014-28491

### Competing Interest Statement

The authors have declared no competing interest.

## References

Arzt, M., Deschamps, J., Schmied, C., Pietzsch, T., Schmidt, D., Tomancak, P., … & Jug, F. (2022). LABKIT: labeling and segmentation toolkit for big image data. Frontiers in computer science, 4, 10.

Cheesbrough, M. (2005). District laboratory practice in tropical countries, part 2. Cambridge university press.

Bayguinov, P. O., Oakley, D. M., Shih, C. C., Geanon, D. J., Joens, M. S., & Fitzpatrick, J. A. (2018). Modern laser scanning confocal microscopy. Curr Protoc Cytom, 85(1), e39. Cold Spring Harb Protoc. (2007). 10x PBS. Cold Spring Harbor Protocols. http://cshprotocols.cshlp.org/content/2007/4/pdb.rec10768.full?text_only=true

Danon, A., Delorme, V., Mailhac, N., & Gallois, P. (2000). Plant programmed cell death: a common way to die. Plant Physiol. Biochem., 38(9), 647–655.

DeZwaan, T. M., Carroll, A. M., Valent, B., & Sweigard, J. A. (1999). Magnaporthe grisea pth11p is a novel plasma membrane protein that mediates appressorium differentiation in response to inductive substrate cues. Plant Cell, 11(10), 2013–2030.

Domínguez, F., & Cejudo, F. J. (2015). Nuclear dismantling events: crucial steps during the execution of plant programmed cell death. Plant Programmed Cell Death, 163–189.

Dugyala, S., Borowicz, P., & Acevedo, M. (2015). Rapid protocol for visualization of rust fungi structures using fluorochrome Uvitex 2B. Plant Methods, 11, 1–8.

Feckler, A., Schrimpf, A., Bundschuh, M., Bärlocher, F., Baudy, P., Cornut, J., & Schulz, R. (2017). Quantitative real-time PCR as a promising tool for the detection and quantification of leaf-associated fungal species–A proof-of-concept using Alatospora pulchella. PLoS One, 12(4), e017463

Gu, Y., & Innes, R. W. (2012). The KEEP ON GOING protein of Arabidopsis regulates intracellular protein trafficking and is degraded during fungal infection. Plant Cell, 24(11), 4717–4730.

Haueisen, J., Möller, M., Eschenbrenner, C. J., Grandaubert, J., Seybold, H., Adamiak, H., & Stukenbrock, E. H. (2019). Highly flexible infection programs in a specialized wheat pathogen. Ecol. Evol., 9(1), 275–294.

Hood, M. E., & Shew, H. D. (1996). Applications of KOH-aniline blue fluorescence in the study of plant-fungal interactions. Phytopathology, 86(7), 704–708.

Jones, K., Kim, D. W., Park, J. S., & Khang, C. H. (2016). Live-cell fluorescence imaging to investigate the dynamics of plant cell death during infection by the rice blast fungus Magnaporthe oryzae. BMC Plant Biol, 16, 1–8.

Kariyawasam, G.K., Richards, J.K., Wyatt, N.A., Running, K.L.D., Xu, S.S., Liu, Z., Borowicz, P., Faris, J.D. and Friesen, T.L. (2022), The *Parastagonospora nodorum* necrotrophic effector SnTox5 targets the wheat gene *Snn5* and facilitates entry into the leaf mesophyll. New Phytol, 233: 409–426. 10.1111/nph.17602

Koladia, V. M., Richards, J. K., Wyatt, N. A., Faris, J. D., Brueggeman, R. S., & Friesen, T. L. (2017). Genetic analysis of virulence in the Pyrenophora teres f. teres population BB25× FGOH04Ptt-21. Fungal genetics and biology, 107, 12–19.

Kumar, A., Karre, S., Dhokane, D., Kage, U., Hukkeri, S., & Kushalappa, A. C. (2015). Real-time quantitative PCR based method for the quantification of fungal biomass to discriminate quantitative resistance in barley and wheat genotypes to fusarium head blight. Journal of Cereal Science, 64, 16–22.

Lagopodi, A. L., Ram, A. F., Lamers, G. E., Punt, P. J., Van den Hondel, C. A., Lugtenberg, B. J., & Bloemberg, G. V. (2002). Novel aspects of tomato root colonization and infection by *Fusarium oxysporum* f. sp. *radicis-lycopersici* revealed by confocal laser scanning microscopic analysis using the green fluorescent protein as a marker. Molecular Plant-Microbe Interactions, 15(2), 172–179.

Liu, Z. H., Friesen, T. L., Rasmussen, J. B., Ali, S., Meinhardt, S. W., & Faris, J. D. (2004). Quantitative trait loci analysis and mapping of seedling resistance to Stagonospora nodorum leaf blotch in wheat. J Phytopathol., 94(10), 1061–1067.

O’Connell, R., Herbert, C., Sreenivasaprasad, S., Khatib, M., Esquerré-Tugayé, M. T., & Dumas, B. (2004). A novel Arabidopsis-Colletotrichum pathosystem for the molecular dissection of plant-fungal interactions. Mol Plant Microbe Interact, 17(3), 272–282.

Redkar, A., Hoser, R., Schilling, L., Zechmann, B., Krzymowska, M., Walbot, V., & Doehlemann, G. (2015). A secreted effector protein of Ustilago maydis guides maize leaf cells to form tumors. Plant Cell, 27(4), 1332–1351.

Shjerve, R. A., Faris, J. D., Brueggeman, R. S., Yan, C., Zhu, Y., Koladia, V., & Friesen, T. L. (2014). Evaluation of a Pyrenophora teres f. teres mapping population reveals multiple independent interactions with a region of barley chromosome 6H. Fungal Genet. Biol, 70, 104–112.

Skipp, R. A., & Deverall, B. J. (1972). Relationships between fungal growth and host changes visible by light microscopy during infection of bean hypocotyls (Phaseolus vulgaris) susceptible and resistant to physiological races of Colletotrichum lindemuthianum. Physiol Plant Pathol, 2(4), 357–374.

Solanki, S., Ameen, G., Borowicz, P., & Brueggeman, R. S. (2019). Shedding light on penetration of cereal host stomata by wheat stem rust using improved methodology. Sci. Rep., 9(1), 1–13.

Spellig, T., Bottin, A., & Kahmann, R. (1996). Green fluorescent protein (GFP) as a new vital marker in the phytopathogenic fungus Ustilago maydis. Mol Genet Genomics, 252, 503–509.

St. Croix, C. M., Shand, S. H., & Watkins, S. C. (2005). Confocal microscopy: comparisons, applications, and problems. Biotechniques, 39(6), S2–S5.

Suzuki, T., Fujikura, K., Higashiyama, T., & Takata, K. (1997). DNA staining for fluorescence and laser confocal microscopy. J.Histochem. Cytochem., 45(1), 49–53.

Tapio, E., & Pohto-Lahdenperä, A. (1991). Scanning electron microscopy of hyphal interaction between Streptomyces griseoviridis and some plant pathogenic fungi. Agric. Food Sci., 63(5), 435–441.

Widengren, J., Mets, Ü., & Rigler, R. (1999). Photodynamic properties of green fluorescent proteins investigated by fluorescence correlation spectroscopy. Chemical Physics, 250(2), 171–186.

