## Supplementary material for "Unleashing the secrets of plant-fungal interactions using a transformation-free confocal staining technique that supports AI-assisted quantitative analysis": Figure S1

A

B

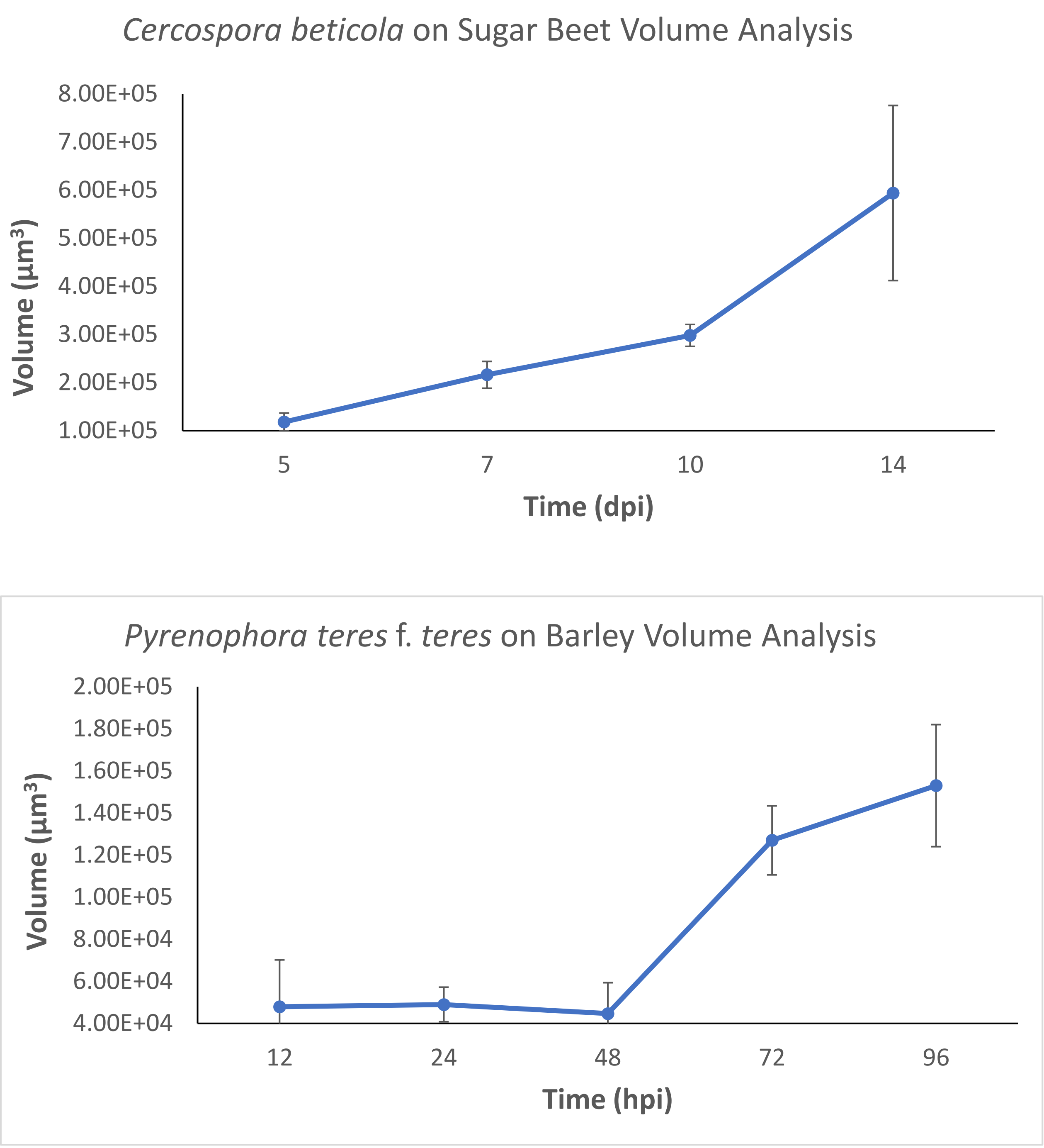
*Cercospora beticola* on Sugar Beet Volume Analysis
Volume (μm³)
8.00E+05
7.00E+05
6.00E+05
5.00E+05
4.00E+05
3.00E+05
2.00E+05
1.00E+05
5
7
10
14
Time (dpi)
*Pyrenophora teres* f. *teres* on Barley Volume Analysis
Volume (μm³)
2.00E+05
1.80E+05
1.60E+05
1.40E+05
1.20E+05
1.00E+05
8.00E+04
6.00E+04
4.00E+04
12
24
48
72
96
Time (hpi)

**Fig. S1** Single replication validations of *Cercospora beticola*-sugar beet and *Pyrenophora teres f. teres*-barley volume analysis

**A**, Fungal biomass of *Cercospora beticola* on sugar beet at the timepoints 5, 7, 10 and 14dpi. Fungal volume begins at $1.18 \times 10^{5}$ $\mu m^{3}$ at five days post inoculations, then increases to $2.16 \times 10^{5}$ $\mu m^{3}$ at seven days post inoculation. A continued increase of $2.98 \times 10^{5}$ $\mu m^{3}$ at ten days is observed followed by a very large increase to $5.94 \times 10^{5}$ $\mu m^{3}$ at 14 days post inoculation. **B**, Fungal biomass of *Pyrenophora teres f. teres* on barley at the timepoints 12, 24, 48, 72 and 96hpi. Fungal volume begins at $4.80 \times 10^{4}$ $\mu m^{3}$ at 12hpi, followed by a slight increase of at $4.90 \times 10^{4}$ $\mu m^{3}$ at 24hpi. A dip in fungal volume is seen at 48hpi, $4.47 \times 10^{4}$, due to a shift in image focus on progressive growth into the mesophyll. A large increase of $1.27 \times 10^{5}$, is observed at 72hpi and is followed by a slight increase of $1.53 \times 10^{5}$ at 96hpi.
